# Feeding rapidly alters microbiome composition and gene transcription in the clownfish gut

**DOI:** 10.1101/440669

**Authors:** D. Joshua Parris, Michael M. Morgan, Frank J. Stewart

## Abstract

**Background:** Diet is a major determinant of intestinal microbiome composition. While studies have evaluated microbiome responses to diet variation, less is understood of how the act of feeding influences the microbiome, independent of diet type. Here, we use the clownfish *Premnas biaculeatus*, a species reared commonly in ornamental marine aquaculture, to test how the diversity, predicted gene content, and gene transcription of the microbiome vary over a two-day diurnal period with a single daily feeding event. This study used fish fed four times daily, once daily, or every three days prior to the diurnal period, allowing us also to test how feeding frequency affected microbiome diversity. The amount of time between feedings had no affect on baseline diversity of the microbiome. In contrast, the act of feeding itself caused a significant short term change in the microbiome, with microbiome diversity, predicted gene content, and gene transcription varying significantly between time points immediately before and 1.5 hours post feeding. Variation was driven by abundance shifts involving exact sequence variants (ESVs), with one ESV identified as *Photobacterium* sp. increasing from <0.5% of sequences immediately pre-feeding to 34% at 1.5 hours post-feeding. Other ESVs from a range of microbial groups also increased dramatically after feeding, with the majority also detected in the food. One ESV identified as *Clostridium perfringens* represented up to 55% of sequences but did not vary significantly over the diurnal period and was not detected in the food. Post-feeding samples were enriched in transcripts and predicted genes for social interactions, cell motility, and coping with foreign DNA, whereas time points farther from feeding were enriched in genes of diverse catabolic and biosynthetic functions. These results confirm feeding as a significant destabilizing force in clownfish intestinal microbiomes, likely due to both input of cells attached to food and stimulation of resident microbes. Microbes such as *Photobacterium* may episodically transition from environmental reservoirs to growth in the gut, likely in association with food particles. This transition may be facilitated by functions for navigating a new environment and interacting with neighboring microbes and host cells. Other taxa, such as *Clostridium*, are comparatively stable intestinal members and less likely to be affected by passing food. Conclusions about microbiome ecology may therefore differ based on when samples were collected relative to the last feeding.

**Importance:** Despite extensive study of intestinal microbiome diversity and the role of diet type in structuring gut microbial communities, we know very little about short-term changes in the intestinal microbiome as a result of feeding alone. Sampling microbiomes over a feeding cycle will allow us to differentiate opportunistic, feeding-responsive microbes from resident, potentially commensal members of the gut community. Also, since feeding has the potential to alter microbiome structure, sampling at different points relative to the last feeding event will likely yield different conclusions about microbiome composition and function. This variation should be addressed in comparative microbiome studies. Our study contributes to knowledge of short-term changes in the gut microbiome associated with feeding events.

## BACKGROUND

The biological significance of host-associated microbiomes is widely recognized, with the intestinal (gut) microbiome in particular now known to play a vital role in host health (1,2). Factors shaping the intestinal microbiome are complex, and include host phylogeny, diet, age, and immune status (3,4). Understanding these factors is essential, as changes in microbiome composition are linked to diverse aspects of host physiology including efficiency of nutrient acquisition, development of the intestine, immune function, and cognition and behavior (2,5,6). However, for most organisms, particularly for non-mammal systems, we lack basic knowledge of how and over what timescale intestinal microbiomes change in response to perturbations, including changes in chemical availability due to feeding.

A lack of knowledge of the dynamics and drivers of microbiome change due to feeding is due partly to the fact that few studies have sampled the intestine over the course of a feeding cycle, which requires sacrificing animals at hourly timescales. Furthermore, interpreting short-term microbiome fluctuations is challenging, as these may also be due partly to host circadian rhythms independent of feeding (7,8), or potentially to changes in feeding frequency (9,10). Zarrinpar et al. (11), for example, compared mice allowed to feed *ad libitum* to mice fed once a day and found that both groups exhibited predictable diurnal oscillations in intestinal taxa, but differed in the number of taxa exhibiting oscillations. In addition, reduced feeding frequency during times when mice were most active was correlated with lower abundance of *Lactobacillus* bacteria, a pattern associated with protection from metabolic disease (12).

The act of feeding may influence microbiome composition and function through diverse mechanisms, independent of diet type. Food intake can introduce new microbes or genes to the intestine (13). Research on probiotic use in gnotobiotic mice shows that microbes from fermented milk products are detected in stool in a matter of days after consumption (14). However, the extent to which the intestinal microbiome is restructured by food-attached microbes in non-model animals has been largely unstudied, although hypothesized to be potentially significant (15). This is surprising, as such restructuring, particularly if transient, could bias conclusions about which microbes live as residents in a stable and potentially beneficial relationship with the host.

Feeding might also stimulate the growth of microbes already present in the intestine (residents). The growth response of residents could be associated with both changes in community structure as well as metabolic cascades linked to food breakdown. Microbes and enzymes specialized for the catabolism of complex carbohydrates might be abundant early in digestion, creating products that can be used for energy by different microbes later in digestion (16,17,18). Such cross-feeding is common in the mammalian intestine and a potential determinant of microbial richness (19,20). Feeding may also promote successional patterns in microbial growth or metabolism independent of metabolite exchange. The mammalian large intestine, for example, is dominated by anaerobic microbes in the mid-lumen, but also contains aerobic or microaerobic taxa, notably along the submucosal surface closer to the oxygenated blood (21,22,23). Similar to what has been shown in chemostat cultures (24), oxygen consumption by aerobes after feeding may precede or promote anaerobic metabolism by other microbes. Indeed, the progression of digestion in insects (from foregut to the hindgut) is characterized by a linear decrease in oxygen concentration (25,26).

The frequency of feeding might also affect the magnitude of microbiome change in response to feeding. Continuous or near-continuous feeding, by grazing animals for example, may maintain relatively constant substrate conditions in the gut as well as a steady stream of food-associated microbes, and therefore a stable assemblage of microbes with slight compositional shifts post-feeding. In contrast, intermittent feeding may promote large compositional changes associated with the transition from a relatively inactive, but stable, “fasting” microbiome to a “bloom” community after feeding. In vertebrate guts, fasting/feeding cycles have been shown to drastically alter the abundance of individual bacterial groups, with fasting associated with higher occurrence of Bacteriodetes (27,28). In addition to altering the magnitude of microbiome change in response to feeding, differences in feeding regime have been shown to affect baseline microbiome composition and metabolite production (7,12). However, the factors driving microbiome differences linked to feeding regime are likely complex, and potentially related to changes in host physiology (29), as well as sampling the microbiome at varying stages in the digestive cycle.

The studies mentioned above, conducted primarily in mammalian models, highlight the need to account for short-term shifts in microbiome composition in comparative studies. However, the extent to which microbiomes of other major animal groups exhibit short-term fluctuation in response to feeding, or other diurnal cues, remains largely uncharacterized. Here, we test how the act of feeding and feeding frequency affect the intestinal microbiome of maroon clownfish (*Premnas biaculeatus*). Fishes are among the most species-rich and ecologically important vertebrate groups, with diverse roles in food webs and as targets of human recreational and commercial interest. Fisheries and aquaculture are multibillion-dollar industries, yet baseline knowledge of the diversity, function, and dynamics of fish microbiomes is absent for most commercially important fish species. *Premnas biaculeatus* in particular is widely bred for the marine ornamental fish trade, is hardy during captivity and in experiments, and represents one of the most abundant and widespread Families (Pomacentridae) in tropical seas. Using this species as a model in feeding experiments, we show large changes in the diversity and presumed metabolic function of the microbiome over a daily feeding cycle but minimal prolonged effect of feeding frequency on baseline microbiome composition. We discuss these results in the context of taxon-specific differences in microbial lifestyle and the potential for feeding events to influence conclusions about microbiome stability and ecology.

## RESULTS

We sampled a cohort of maroon clownfish (*Premnas biaculeatus*) at five time points per day over a two-day period with one feeding event per day (at 1100). Before sampling occurred, fish were divided among three treatment groups differing based on whether they were fed four times daily (4X, n=29), once daily (1X, n=29), or once every three days (0.33X, n=30) over the month prior to the two-day sampling period. Three fish per treatment group were sacrificed immediately pre-feeding (1100, n=17), 1.5 hours post-feeding (1230), 3 hours post-feeding (1400), 5 hours post-feeding (1600, n=17), and 9 hours post-feeding (2000, n=17) on each of the two days of the sampling period. Thus, each time point (1100, 1230, 1400, 1600, and 2000) was represented by 18 individuals (3 replicates per daily time point X 2 days of sampling X 3 treatment groups; for some time points, we obtained data from 17 rather than 18 individuals due to sample/fish loss). These samples were used for analysis of the intestinal microbiome along with samples of the water and food (see Methods for full experimental and sampling details).

## Microbiome taxonomic composition

We recovered 6,732,587 16S rRNA gene amplicon sequences (range: 706-239,836 per sample; average: 64,120) and 2748 unique exact sequence variants (ESVs) across all samples of fish, tank water, and food (average: 288 ESVs per sample) (Table S1). Microbiome composition in water and food samples differed significantly from that of the fish intestine (PCoA, ANOVA, P<0.05; Figure S1). More than 90% of sequences in the water (across all replicates) were identified as belonging to the bacterial Families Flavobacteriaceae (Bacteroidetes), Methylophilaceae (Alphaproteobacteria), or Rhodobacteraceae (Alphaproteobacteria). Food microbiomes were dominated (~70%) by sequences of the Phormidiaceae and Streptophyta (Cyanobacteria), Pseudoalteromonadaceae (Gammaproteobacteria), and Rickettsiales (Alphaproteobacteria) (Figures S1-S2). In contrast, fish microbiomes were composed primarily (~70%) of sequences of the Families Clostridiaceae (Firmicutes), Mycobacteriaceae (Actinobacteria), and Vibrionaceae (Gammaproteobacteria; Figure S1). Many groups, notably Vibrionaceae, were detected in both the intestine and food samples (see below). Alpha diversity (Shannon index) did not differ significantly among water, food, and fish samples (fish vs. water vs. food, all samples combined).

Fish microbiome composition and alpha diversity during the two-day diel sampling period did not differ among samples grouped based on feeding treatment (4X, 1X, or 0.33X daily feedings in the month prior to sampling; Figure S3). Furthermore, no ESVs were detected as differentially abundant among feeding groups. Similarly, no differences were observed when analyzing only samples from the first time point (pre-feeding, 1100) on day 1 of the diel sampling, prior to the synchronization of feeding schedules for the diel sampling, and presumably the point at which the intestinal communities of different groups would be most affected by prior feeding regime.

In contrast, microbiomes varied significantly among samples grouped based on time of day during the two-day diel sampling period, irrespective of feeding frequency treatment in the prior 30 days (Figure 1A, Figures S4-S5). Microbiomes generally partitioned into two clusters, one including nearly all samples from the two time points immediately post-feeding (1230, 1400; “fed” in Figure 1A), and another including the two time points most separated from the feeding event (1100, 2000; “unfed”). Microbiomes at the intermediate time point (1600, 5 hours post-feed) were more variable, with replicates falling within both the fed and unfed clusters. Consistent with these clustering patterns, random forest analysis based on 698 ESVs (best model) showed that microbiomes from the 1100 and 1230 time points could be correctly classified to those time points with 100% accuracy, whereas ESV composition was less predictive of later time points post-feeding (Figure 2).

**Figure 1.**
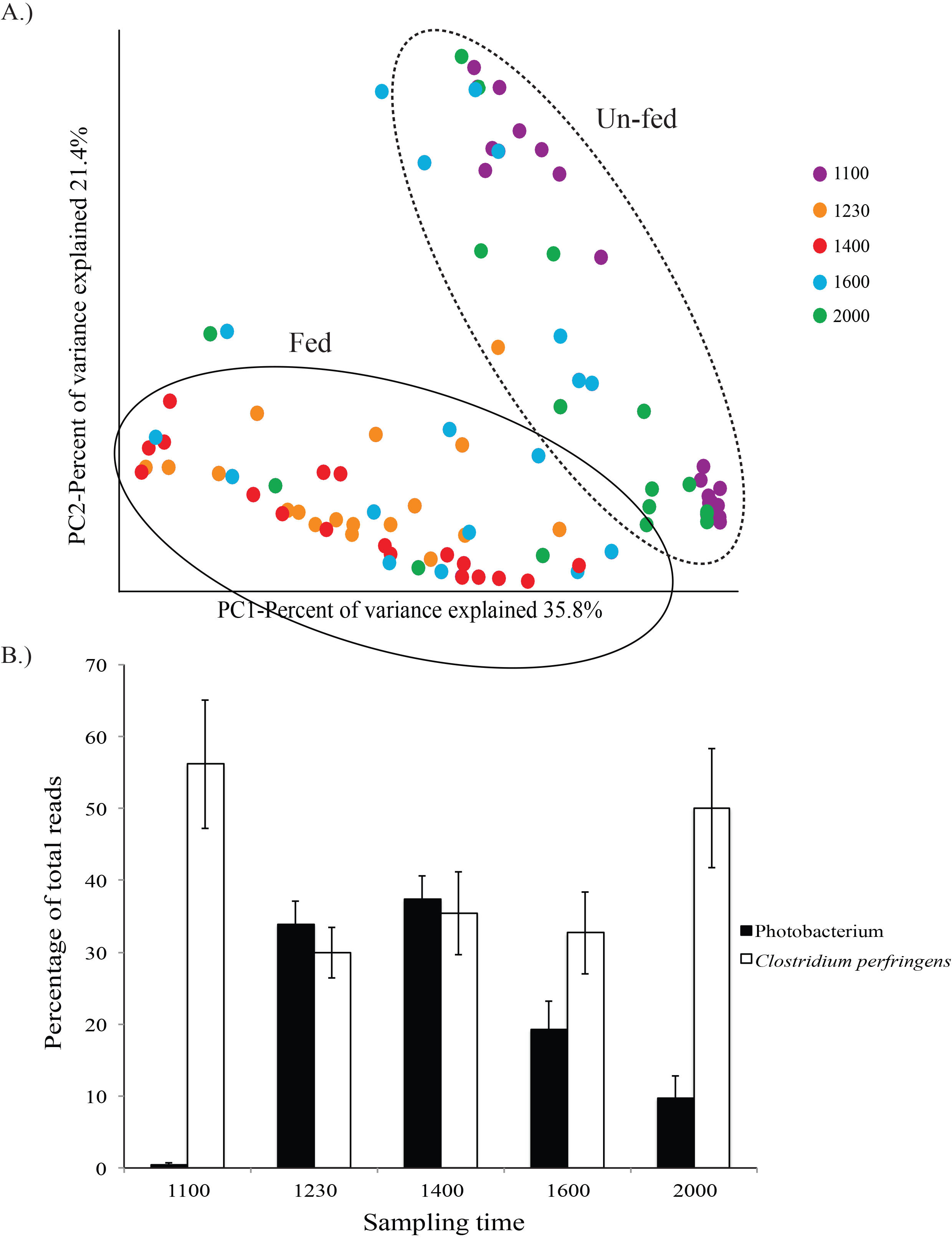
Intestinal microbiome composition varies over a feeding cycle A.) Principal coordinate analysis showing significant microbiome clustering by time of sampling (ANOVA, P<0.05). Distance was based on Bray-Curtis similarities with all samples rarefied to 20,562 sequences. Samples within the dashed circle were primarily from the unfed time points while samples within the solid circle were from the fed time points. B.) Relative abundance of the two most common ESVs (exact sequence variants) varies over a feeding cycle (n=17 at 1100 and 2000, n=18 at 1230, 1400, and 1600). ANCOM analysis identified the *Photobacterium* ESV, but not the *Clostridium* ESV, as varying significantly over the sampling period (P<0.05).

**Figure 2.**
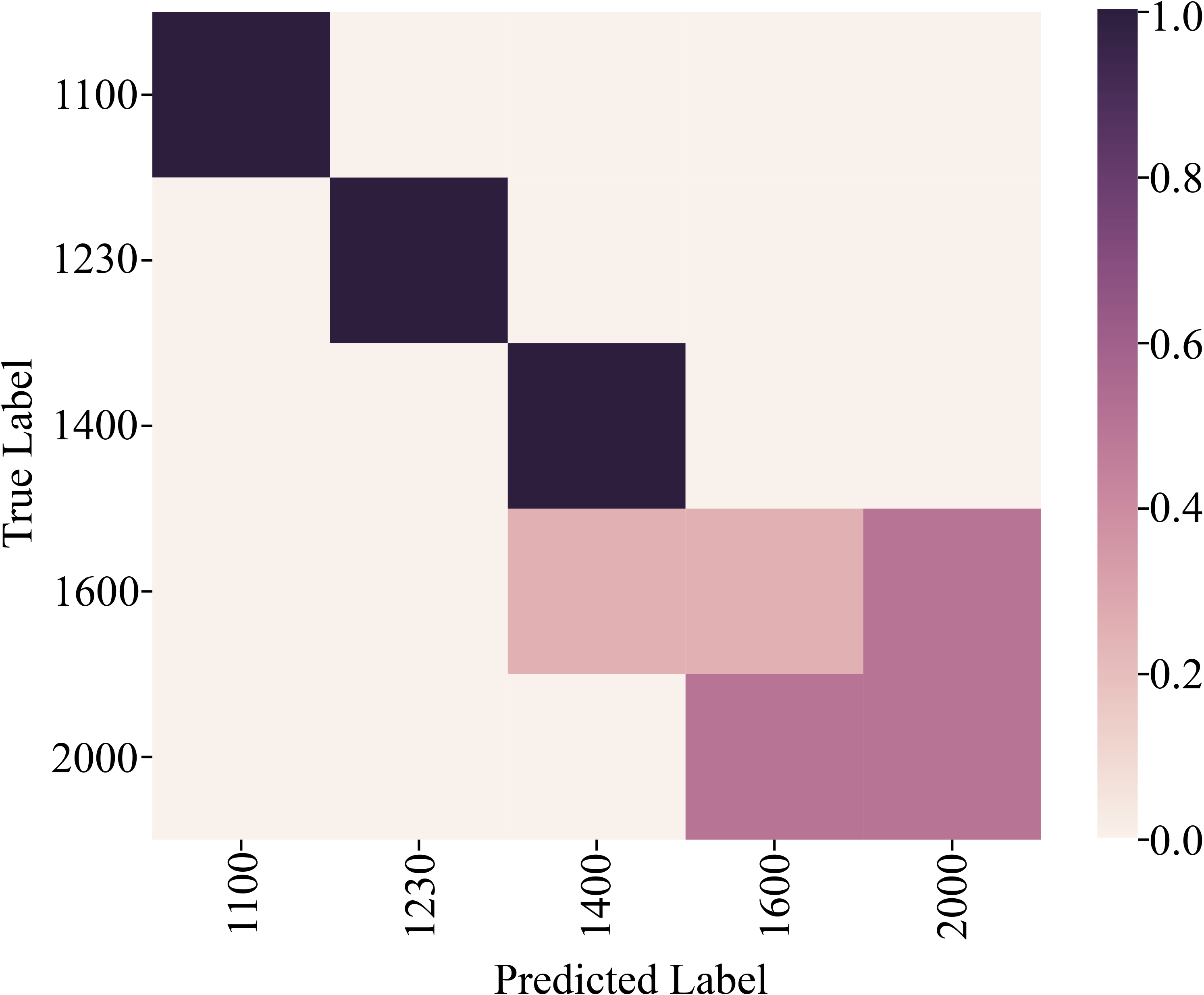
Samples can be accurately classified to sampling time based on community composition. Random forest model showing the frequency at which microbiome samples from a given time point are correctly assigned based on taxonomic composition. This model included 698 unique sequence features (best model) and had an overall accuracy of 72% (n=17 fish at 1100 and 2000 and18 fish at 1230, 1400, and 1600).

Average alpha diversity (Shannon index) also varied significantly based on sampling time, being lowest immediately pre-feeding at 1100, increasing by over 2-fold by 1.5 hours post-feeding (1230), and then decreasing steadily thereafter as intestinal content cleared (Figure 3). Shannon diversity at 1230 was significantly higher compared to every other time point, excluding 1600; diversity at 2000 was significantly lower compared to every other time point, excluding 1100 (Kruskal Wallis pairwise, corrected p-value <0.05).

**Figure 3.**
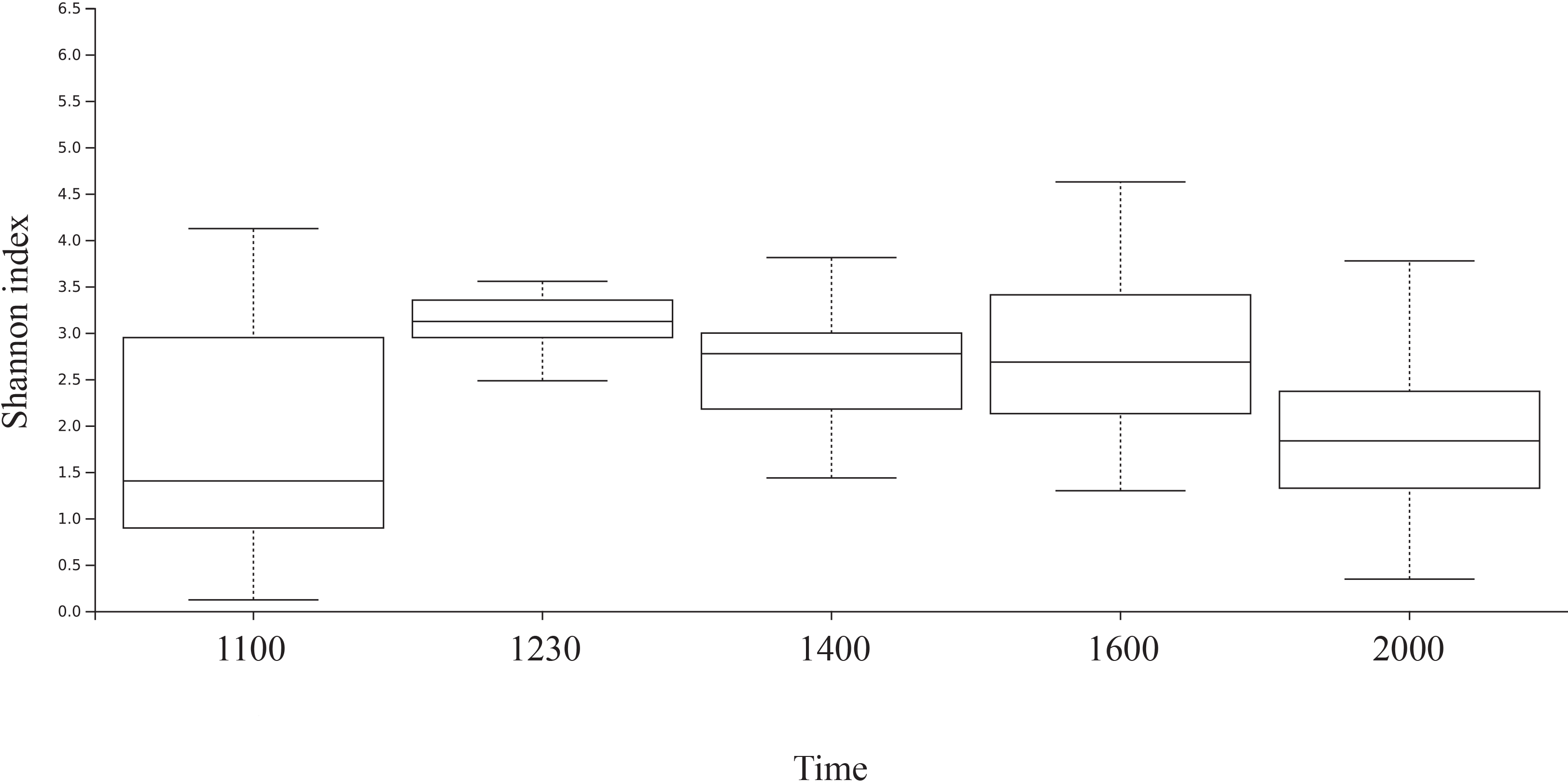
Average Shannon diversity varies according to time of sampling. Shannon diversity at 1230 was significantly higher compared to every other time point, excluding 1600; diversity at 2000 was significantly lower compared to every other time point, excluding 1100 (Kruskal Wallis pairwise, corrected p-value <0.05, n=17 fish at 1100 and 2000 and 18 fish at 1230, 1400, and 1600).

Diverse microbial groups fluctuated in abundance over the two-day sampling (Figure S4, Table S2). The most abundant ESV in the dataset was classified as *Clostridium perfringens* (Firmicutes). This ESV comprised 50-56% of all amplicons at unfed time points (1100, 2000) and 29-35% at the intermediate, fed time points (Figure 1B); however, this variation was not statistically significant (P>0.05) based on analysis of composition of microbiomes (ANCOM, Mandal et al. 2015). ANCOM analysis, however, identified 57 ESVs whose abundance varied significantly based on time (Table S2). Almost all of these (55 of 57) increased markedly in representation from pre-feeding (1100) to 1.5 hours post-feeding (1230). Remarkably, an ESV identified as *Photobacterium* sp. increased from <0.5% of amplicons in pre-feeding samples to 34% of amplicons during this period (Figure 1B; Table S2). Its abundance peaked at 1400, and then decreased sharply over the remaining time points. Other significantly varying ESVs that followed the same trend, but were less abundant, included unclassified members of the Vibrionaceae and diverse members of the phylum Firmicutes; roughly one-third of the significantly varying ESVs were identified as bacteria of the class Bacilli (Firmicutes), with all but one of these showing multi-fold increases in representation from the 1100 to 1230 time point (Table S2). A smaller number of significantly varying ESVs showed the opposite trend, decreasing in representation after feeding. These included ESVs of the Firmicutes genus Streptococcus and unclassified members of the order gammaproteobacterial order Alteromonadales (Table S2).

Of the 57 significantly varying ESVs, all but one were also detected in the food, albeit at relatively low proportional abundance (Table S2). Food-associated ESVs included the intestinally abundant *Photobacterium* sp. ESV, which comprised ~2% of food sequences. In contrast, the most dominant food-associated sequences (>10%), belonging to the Cyanobacteria (Figure S2), were not among the significantly varying ESVs in the intestine and contributed negligibly to the intestinal dataset.

## Predicted metagenome content

Metagenome prediction based on 16S rRNA gene profiles identified 329 gene categories. Of these, none were differentially represented among the three feeding frequency treatments, based on samples collected at the first 1100 time point of the diel-sampling period. In contrast, 111 functional gene categories were predicted to vary in abundance between unfed (1100 and 2000) and fed (1230, 1400, 1600) time points (DESEQ2, adjusted p<0.05, Table S3). Functions predicted to be enriched in the fed state included those associated with bacteria-bacteria or bacteria-host interactions, including bacterial invasion of epithelial cells, infection by *Vibrio*, secretion, motility, and chemotaxis. In contrast, unfed time points were dominated by anabolic and catabolic functions (Figure 4, Table S3). Of the top 50 predicted functional categories that were significantly enriched in the unfed time points, 76% (38/50) were classified broadly as metabolism, degradation, or biosynthesis, including those for amino acid metabolism, fatty acid metabolism, secondary bile acid production, sphingolipid biosynthesis, and the degradation of a wide range of organic compounds. In comparison, metabolism, degradation, or biosynthesis-associated categories represented only 18% of those enriched in the fed datasets.

**Figure 4.**
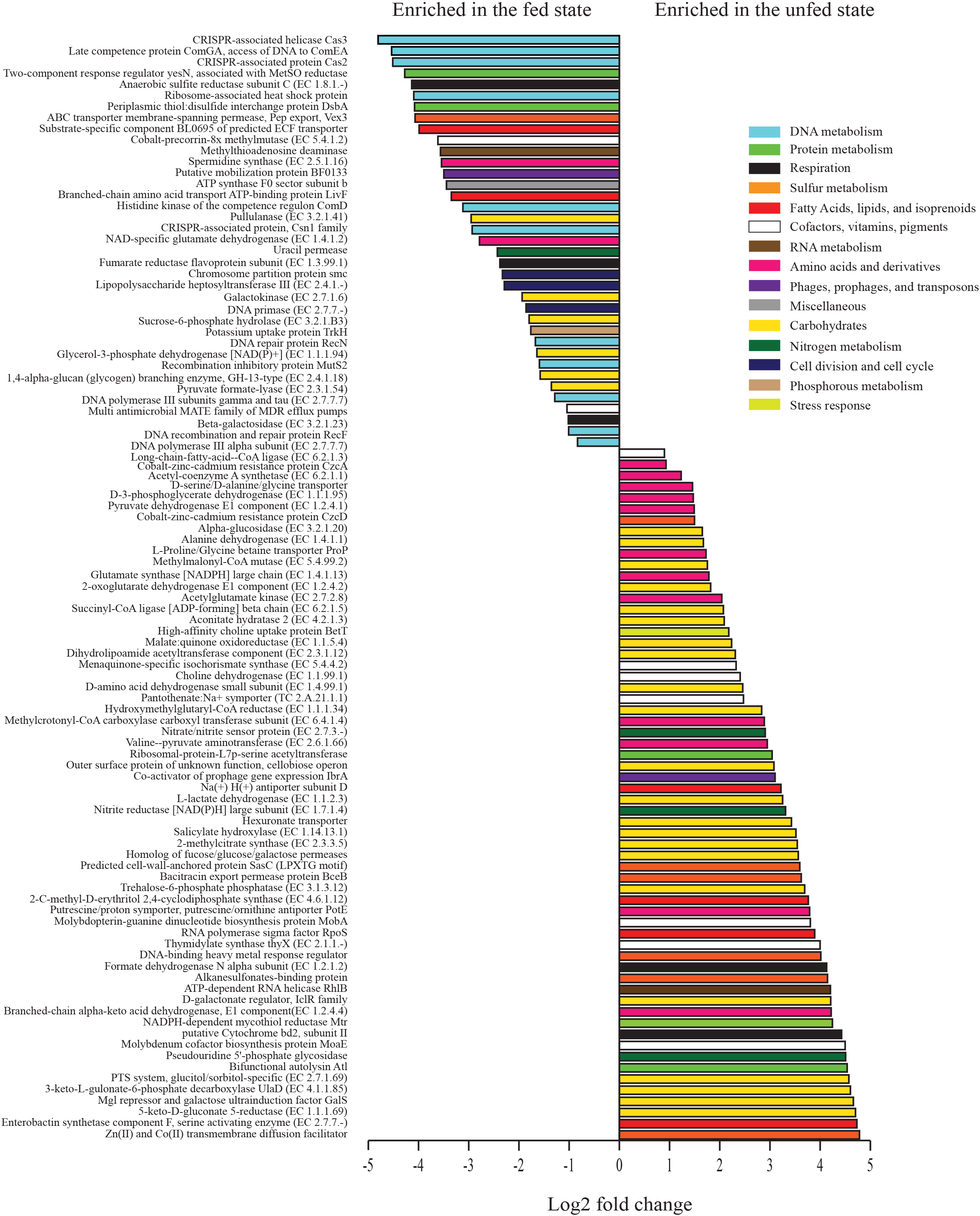
Predicted functional gene categories vary significantly between unfed (1100 and 2000) and fed (1230, 1400, 1600) time points with fed time points enriched in disease-associated pathways and unfed time points enriched in diverse metabolic pathways. Metagenomes were predicted from amplicon data using PICRUST and the 3^rd^ hierarchical level of KEGG. Differential abundance was evaluated using DESEQ2 in R with gene categories having an adjusted p-value <0.05 shown here. Colors represent larger subsystem categories in KEGG. Only the top 99 of 111 significant pathways are plotted (n=17 fish at 1100 and 2000 and 18 fish at 1230, 1400, and 1600)..

## Differential transcription

Metatranscriptome sequencing yielded 2,188,905 non-host, mRNA reads; per-sample counts ranged from 58,611 to 681,223 (Table S4). Of these, 166,307 reads (across all samples) were classified as bacterial and had functional matches in the SEED database (a constantly updated repository for genomic sequence information; Overbeek et al. 2005), representing 1,063 pathways (second level of the SEED classification). Within these SEED pathways, a total of 267 genes showed differential transcription between fed and unfed samples (p<0.05, DESeq, Table S5). Of the top 100 most differentially transcribed genes (based on adjusted p-value), 62 were at higher abundance in the unfed state (Figure 5). Consistent with the metagenome predictions, unfed transcriptomes were enriched in functions associated with metabolism, notably carbohydrate and amino acid metabolism. Over one third of the genes most enriched in the unfed transcriptomes were associated with diverse steps of carbohydrate utilization (compared to 16% of the fed-enriched genes), including several associated with pyruvate metabolism and the citric acid cycle (EC 1.2.4.2, EC 6.2.1.5, EC 4.2.1.3, EC 1.1.5.4, EC 2.3.1.12, EC 1.1.2.3, EC 2.3.3.5), fermentation or the metabolism of fermentation intermediates (EC 1.1.2.3, EC 2.3.3.5), and the degradation of cellulose or other complex organic molecules (EC 1.14.13.1, EC 3.1.3.12, EC 1.1.1.69). In contrast, fed state transcriptomes were enriched in genes of DNA metabolism. This included several genes associated with CRISPR defense systems to cope with foreign DNA, as well as genetic elements associated with recombination (Figure 5).

**Figure 5.**
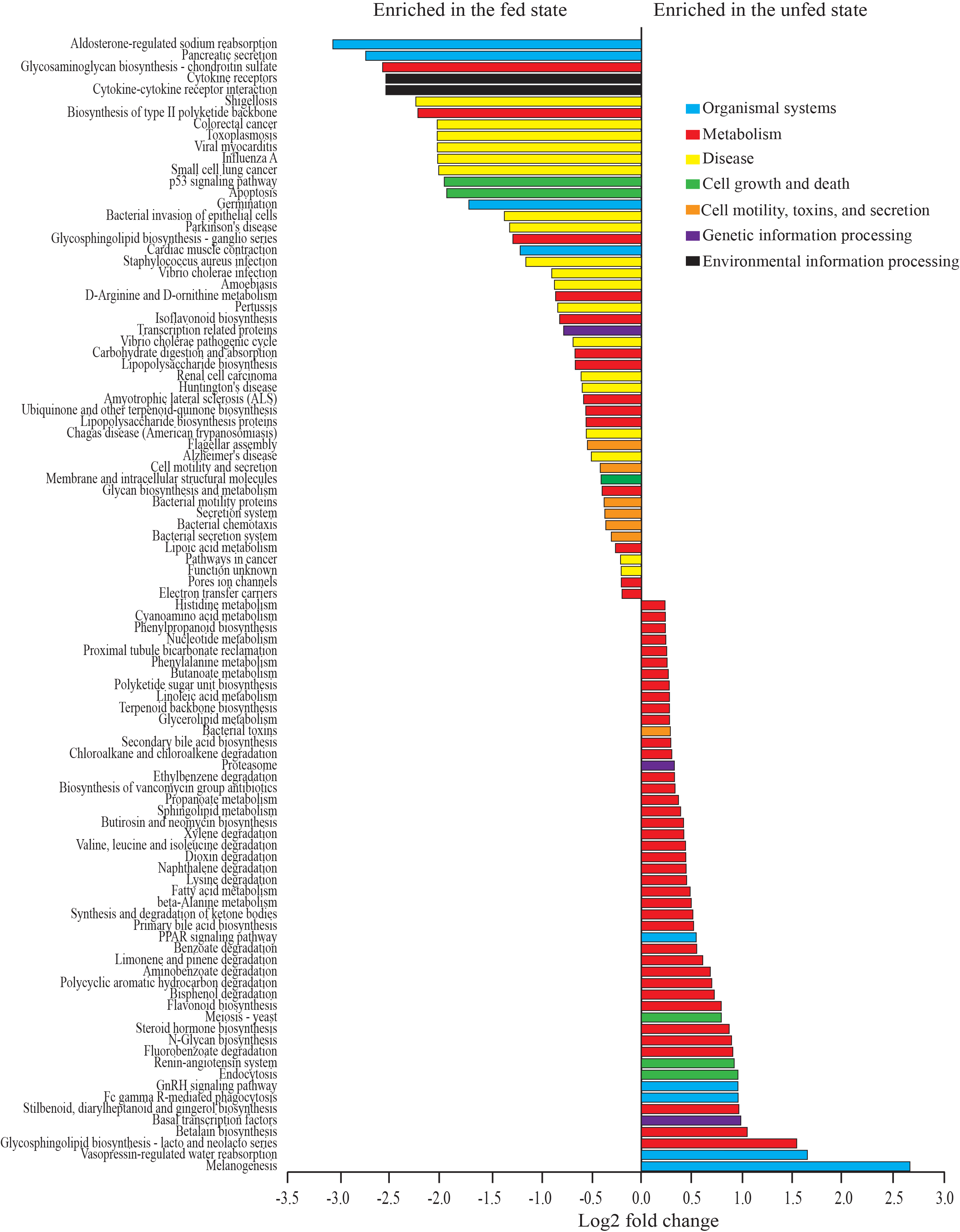
Gene expression varies significantly over a diumal feeding cycle. Top SEED functional genes (by adjusted p-value) showing differential expression between fed and unfed states in transcriptomic data (1100 vs 1230 time points, n=5 for each). Differential abundance was tested with DESEQ2 in R and all functional categories shown vary significantly (p<0.05). Colors represent larger subsystem categories in SEED.

## DISCUSSION

We used a popular and commonly bred marine aquarium species, the clownfish *Premnas biaculeatus*, as a model to explore how feeding events and feeding frequency alter the gut microbiome. Quantifying microbiome changes over the short time frame of a feeding and digestive cycle is critical for evaluating if microbiome studies should standardize sampling around feeding schedules, characterizing microbiome stability, distinguishing resident microbiome members from transient ones, and determining the extent to which the microbiome can be shaped by changes in feeding strategy independent of diet type.

In our experiment, gut microbiome composition and predicted metabolic function varied significantly over a 24-hour cycle (evaluated over two days). This cycle included one feeding event per day, and the most substantial changes in microbiome composition were evident when samples were grouped in relation to this event (“fed” vs “unfed”). Microbiome alpha diversity also spiked after feeding, suggesting that new microbes were introduced via food or that feeding changed the growth dynamics of resident microbes, or both. These patterns implicate feeding, rather than other host diel rhythms, as the primary driver of microbiome change over short (hourly) timescales in our experiment. In contrast, microbiome communities, when analyzed at the start of the two-day diel sampling period did not group based on the frequency of feeding over the prior 30 days, suggesting that prior feeding frequency did not have a lasting restructuring effect on the microbial community.

Variation in microbiome composition over the diel period was driven largely by individual sequence variants, notably a member of the gammaproteobacterial genus *Photobacterium*. Its abundance in the intestine increased from near zero immediately before feeding to over one third of all sequences 1.5 hours post-feeding. This ESV was present in the food and therefore its post-feeding increase may be due to the arrival of new cells in the intestine, a process consistent with the observed post-feeding increase in alpha diversity. However, feeding also may have stimulated growth of cells already in the intestine. Indeed, *Photobacterium* species are common in fish microbiomes (30,31). Some species are pathogens, while others play mutualistic or commensal roles (32,33). Members of the genus are typically facultatively aerobic chemoorganotrophs, motile via flagella, and employ diverse mechanisms for extracellular signaling and host interaction, including multiple virulence factors (34). *Photobacterium* genomes often contain a high number of rRNA operons (often >10; 35), which biases estimates of the proportional abundance of this genus using 16S rRNA gene data. Nonetheless, the magnitude of change in *Photobacterium* sequence abundance after feeding is significant and implicates this taxon as an opportunistic member of the gut whose abundance and activity are closely linked to food availability. Indeed, the genus has been suggested to play a role in fish digestion, potentially by aiding the breakdown of chitin (36,37,38). Follow-up experiments that vary the chitin content of the diet could be used to test a linkage between *Photobacterium* population oscillations and chitin metabolism.

Other ESVs also fluctuated dramatically over the diel period but were less abundant (typically <0.5%). Like *Photobacterium*, the majority of these bacteria were also detected at low abundance in the food, barely detectable in pre-feeding samples, and spiked in representation immediately post-feeding (1230). Many of these taxa were members of the phylum Firmicutes, including diverse genera of lactic acid bacteria (LAB; order Lactobacillales) such as *Lactobacillus*, *Vagococcus*, *Leuconostoc*, and *Streptococcus* (Table S2). LAB are common in vertebrate gut microbiomes, including of fishes in which *Lactobacillus* diversity in particular has been shown to be highly responsive to diet shifts (39,40). In mammalian systems, LAB have been shown to vary in abundance during feeding and non-feeding phases, although the nature of this variation was taxon-specific and varied depended on the timing (frequency) of feeding and also on diet type (11). LAB, and Firmicutes in general, are thought to be important to host carbohydrate metabolism through fermentation and have been associated with efficient dietary energy harvest (41,42,43); we hypothesize that LAB are likely playing a similar role in fermentation and energy extraction in the clownfish gut, although the specific dietary compounds supporting LAB catabolism in this system remain to be identified. Our findings corroborate prior evidence suggesting that LAB are among the most responsive to feeding, with many taxa showing a positive response, a factor that may also contribute to their role in energy extraction from food.

The post-feeding spike in abundance of *Photobacterium* and other diverse ESVs, coupled with their detection in the food, implicate microbial attachment to food as a major determinant of intestinal microbiome composition. However, other taxonomic groups that were much more abundant in the food (e.g., *Pseudoalteromonas*, diverse Cyanobacteria; Figure S4) were not among those showing significant diurnal variation in the intestine microbiome. This suggests that the observed post-feeding spikes by certain intestinal microbes were not due exclusively to the influx of dead cells or DNA. Rather, microbes such as *Photobacterium* or LAB may be adept at both surviving passage through the stomach and exploiting the intestinal environment for growth, at least in the short term. It is also possible that these taxa are taking advantage of metabolites produced by other microbes or that the introduction of food provides them a competitive advantage by altering physical conditions or promoting growth-limiting factors like phages. Further work is needed to assess how the abundance and metabolism of these feeding-responsive taxa may vary with diet, including in omnivorous hosts such as *Premnas biaculeatus*, and to what extent these taxa persist in the intestine without regular input of new food-associated cells.

Other taxa, some of which were highly abundant, exhibited less dramatic fluctuation over the diel period and were not detected in the food. An ESV most closely related to the Firmicutes bacterium *Clostridium perfringens* dominated (>50%) the intestinal microbiomes in the unfed stages and also remained abundant after feeding. *Clostridium* bacteria are common in the intestine of fishes (31,44,45) and other vertebrates, including in humans where this genus has been associated with mucus scavenging (46) and in mice where *Clostridium* has been shown to vary over a daily cycle (7, 11). These diverse fermenters are some of the first taxa to colonize human infants, are believed to localize to particular epithelial cells in the colon, and are important to colonic health by producing butyrate as an energy source for colonocytes (47). In herbivorous fishes, *Clostridium* species have been associated with potentially beneficial roles in vitamin and fatty acid synthesis (48) and the production of metabolic enzymes for catabolism (49). Species such as *C. perfringens* are common food-born pathogens in humans and have been found previously in diverse fishes (e.g.,50,51). In our study, the proportional representation of *C. perfringens* was undoubtedly influenced by swings in taxa such as *Photobacterium* that increased rapidly after feeding (and vice versa). However, the overall high representation of *C. perfringens* at both fed and unfed time points suggests that this taxon may be physically associated with the intestinal lining, rather than the transitory stool. Association with the intestinal mucosal epithelium would be consistent with the mucolytic capabilities observed previously in *Clostridium* (52) and may suggest *Clostridium* as a comparatively persistent microbiome member across changes in diet or food availability.

Significant changes in community composition over the feeding cycle coincided with differences in predicted gene content and gene transcription in the fed and unfed states. The results were generally consistent between analyses, with time points immediately after feeding enriched in pathways involved in bacterial secretion systems, pathogen interaction with hosts, cell motility, and coping with foreign DNA (e.g., CRISPR). In contrast, unfed time points were enriched in transcripts and predicted genes of diverse metabolic processes, particularly those involved in the catabolism of diverse organic substrates, including through fermentation, suggesting these periods as important for microbial degradation of dietary compounds. This enrichment of genes of fermentative carbohydrate and amino acid metabolism is likely linked to *Clostridia*, the dominant taxonomic group in the unfed state (Figure 1b) and one known to play diverse fermentative roles in other gut systems. These patterns are undoubtedly influenced by the dramatic fluctuations in the feeding-responsive members of the community, primarily *Photobacterium*, but likely also the unclassified members of the Vibrionaceae that increased during fed time points (Table S2). Indeed, comparative analysis of *Photobacterium* genomes has revealed high numbers of CRISPR arrays, prophage sequences, and genomic islands, suggesting that phage infection and gene mobilization may be common in this genus (53); other Vibrionaceae genomes share similar features (54,55).

These results suggest that the period shortly after feeding may be a time of increased bacteria-bacteria and bacteria-host interaction. The movement of food into the intestine may stimulate bacteria living in association with the host mucus layer to mobilize and attach to food particles. These early responders therefore may be those cells best equipped to navigate a dynamic and spatially structured environment. Some of these are likely already resident in the intestine, although perhaps at low proportional abundance. However, our results suggest that feeding also serves as the major inoculation event through which new cells enter the system and potentially try to establish residence, a process that would presumably also be characterized by social interactions, attachment, and motility.

## CONCLUSIONS

These results confirm that feeding is a major restructuring force in intestinal microbiomes over a short timeframe (hours). This restructuring involves swings in proportional abundance that differ among microbial types, likely due to differences in metabolic and spatial niche (for example, attachment to food versus residence in an intestinal epithelial biofilm), and potentially also interactions among neighboring microbes. The patterns reported here identify taxa to target for comparisons of how opportunistic, feeding-responsive microbes of the intestine differ ecologically from more persistent, and potentially commensal, members. The large post-feeding changes involving food-associated microbes indicate high connectivity between external and intestinal microbial pools. Animals rarely if ever consume sterile food, even in captivity, and also exhibit significant variation in feeding schedule. This variation should be addressed in comparative microbiome studies, particularly those involving wild animals or small numbers of replicates, but also those focused on model systems (e.g., gnotobiotic mice). Indeed, a growing body of evidence, including from this study, suggests that the vertebrate gut microbiome can exhibit significant short-term fluctuation. Sampling a microbiome at different points relative to the last feeding event will therefore likely yield different conclusions about microbiome composition and function.

## METHODS

### Feeding experiments

A single cohort of 120, six-month old maroon clownfish (*Premnas biaculeatus*) was obtained from Sustainable Aquatics (Jefferson City, TN) and allowed to acclimate for two weeks in an artificial seawater system at Georgia Tech. All fish were fed a 0.8 mm dry pellet composite of krill meal, fish meal, squid meal, wheat gluten, potato starch, fish oil, spirulina, astaxanthin, and garlic oil produced by Sustainable Aquatics. During acclimation, fish were fed once daily at 1100. Following acclimation, fish were equally divided into 12 identical, 10-gallon tanks, all of which were connected to the same recirculating water system. Individual tanks were randomly assigned to one of three treatment groups based on feeding frequency: a 4x treatment with feeding of 2.1 mg of food per fish 4 times daily at 0800, 1100, 1600, and 2000; a 1x treatment with feeding of 8.4 mg of food per fish once daily at 1100; and a 0.33x treatment with feeding of 25 mg of food per fish once every 3 days at 1100. While we could not verify the actual amount of food eaten per fish, the amount of food applied per feeding in the 0.33x treatment was consumed entirely (the food was buoyant, allowing us to track consumption), indicating that this amount did not exceed the maximum clearance rate of each fish group. The feeding treatments were administered for 30 days and water quality was monitored daily. Samples of the tank water microbiome were collected from each treatment at the midpoint and end of the experiment by filtering 1 L of water onto 0.2 μm 25 mm disc filters (n=3 per time point per treatment). Samples (n=3) to analyze microbes in the food were collected at the end of the experiment. A subset of fish (n=6) were sacrificed before initiating feeding treatments and at day 15 (one from each treatment) during the experiment by submerging individuals in a sterile water bath containing MS-222. After euthanization, a ventral cut was made on each fish to expose the gut cavity to preservative, and each fish was then preserved in RNA/DNA stabilizing buffer (25 mM sodium citrate, 10 mM EDTA, 5.3M Ammonium sulfate, pH 5.2) and stored frozen. After 30 days, the remaining fish from each treatment group were sacrificed at various daily time points over a two day period (3 treatment groups X 3 replicates per daily time point X 2 days of sampling each time point = 18 total individuals sacrificed per time point): immediately pre-feeding (1100, n=17), 1.5 hours post-feeding (1230, n=18), three hours post-feeding (1400, n=18), 5 hours post-feeding (1600, n=17), and nine hours post-feeding (2000, n=17). (For some time points, we obtained data from 17 rather than 18 individuals due to sample/fish loss) During this two-day sacrifice period, all fish were fed only once per day at 1100 regardless of original feeding frequency treatment. Fish were euthanized and preserved as described above. Prior to DNA/RNA extraction, whole intestines were removed from each fish via sterile dissection.

### DNA/RNA extractions and amplicon generation

Total DNA and RNA was extracted from intestinal contents and water/food samples using the Mobio PowerMicrobiome® RNA Isolation kit. Total extracts were split into DNA and RNA pools and treated with either DNase or RNase respectively. Before beginning the kit protocol for fish samples, a longitudinal cut was made along the length of the previously dissected intestine. The intestine and associated contents were placed inside a bead-beating tube provided with the kit, vortexed for 5-10 seconds, and the remaining intact intestine was removed and discarded to minimize host signal. The remainder of the extraction followed standard procedures for the Mobio PowerMicrobiome® RNA Isolation kit.

Illumina sequencing of 16S rRNA gene amplicons was used to assess microbiome community composition in a subset of experimental fish (90 fish total: 6 per time point * 5 time points * 3 treatments). Amplicons were synthesized using Platinum® PCR SuperMix (Life Technologies) with primers F515 and R806 spanning the V3-V4 region of the 16S rRNA gene (56). Forward and reverse primers were modified to include Illumina sequencing adapters according to Kozich et al. (57) and barcoded by sample to maintain integrity of biological replicates. Approximately 5 ng of starting DNA was used as template for each PCR reaction. Negative controls using sterile water were included with each set of PCR reactions. Amplification was performed using denaturation at 94°C (3 min), followed by 30 cycles of denaturation at 94°C (45 sec), primer annealing at 55°C (45 sec), primer extension at 72°C (90 sec), and a final extension at 72°C for 10 min. Amplicons were verified using gel electrophoresis, purified using Diffinity RapidTip2 PCR purification tips (Diffinity Genomics, NY), and quantitated fluorometrically using the Qubit (Life Technologies). Barcoded amplicons from all samples were pooled at equimolar concentrations and sequenced on an Illumina MiSeq using a 500 cycle kit with 10% PhiX added to increase sequence diversity.

### Amplicon sequence analysis

Amplicon sequence data were sorted by sample according to barcode, quality-controlled (removed bases <Q30 and sequences less than 150 base pairs), and clustered into exact sequence variants (ESVs) using Quantitative Insights Into Microbial Ecology (QIIME2, version 2018.2) with plugins demux and deblur (58). Sequences were summarized using the feature-table function and aligned, and representative sequences from each cluster were arranged in a phylogenetic tree using FastTree. The feature table with water and food samples included was rarified to 10,321 sequences per sample and alpha and beta diversity explored using the q2-diversity plugin. Alpha diversity among sample groupings was compared using the Shannon Diversity metric, and beta diversity was compared using Bray-Curtis distance. A second feature table was created to include only experimental fish samples and was rarefied to 20,562 sequences per sample. This table excluded 1 sample from the 1100 time point and one sample from the 2000 time point due to low sequence yield. Taxonomy was assigned to sequence clusters using a pre-trained Naive Bayes classifier and the q2-feature-classifier plugin. This classifier was trained on the Greengenes 13_8 99% OTUs database which included a 250 base pair segment from the V4 region of the 16S. Differentially abundant taxa (grouped at the genus level) between sample groupings (feeding frequency, time of sampling) were identified using ANCOM (59). The ability to predict sample groupings based on taxonomic composition was assessed using the q2-sample-classifier function and a random forest model for both feeding frequency and time of sampling.

Predicted metagenomes were constructed from amplicon data using Phylogenetic Investigation of Communities by Reconstruction of Unobserved States (PICRUSt, 60). This analysis was based on an OTU table generated using QIIME1 with the pick_closed_reference_otus.py command, OTUs clustered at 97% sequence similarity, and Greengenes version 13_8. QIIME1 was used at this step since the output OTU table is compatible with PICRUSt. The OTU table was normalized by dividing each OTU by the known/predicted 16S rRNA gene copy number with the normalize_by_copy_number.py command in PICRUSt. Functional predictions based on KEGG categories were made via the predict_metagenomes.py command and collapsed at the 3^rd^ hierarchical level using the categorize_by_function.py command. Predicted functional categories that differed in abundance between feeding treatments and sampling time points were identified using DESeq2 in R.

For ease of comparison and because taxonomic analysis showed high similarity among post-feeding time points, samples were grouped into “fed” (1230, 1400, and 1600) and “unfed” (1100 and 2000) categories. The “unfed” grouping is presumed to include time points at which most of the ingested food has left the stomach; however, it is estimated to take ~36 hours for food to completely pass through the clownfish intestine (61). Therefore, it is likely that all samples contained some amount of food regardless of sampling time.

### RNA sequencing and analysis

The amplicon-based analyses (above) suggested a large difference in composition and predicted microbiome function between samples in the fed and unfed states, primarily at the time points before and shortly after feeding. We therefore chose samples on day 1 from the fed (1230 time point, n=5) and unfed (1100 time point, n=5) state as focal points to analyze transcription of metabolic genes in response to feeding. To generate mRNA data, DNA was first removed from an aliquot of each RNA extract using the TURBO DNA-*free*™ kit (Invitrogen). We next attempted to enrich samples for non-host, non-ribosomal RNA using the MICROBEnrich™ kit (Ambion) and the Ribo-Zero rRNA removal kit (Illumina) following recommended procedures. Barcoded cDNA libraries were prepared from microbial mRNA-enriched RNA using the ScriptSeq kit by Illumina and sequenced (250 × 250 bp) on one lane of an Illumina HiSeq 2500 flowcell in Rapid mode.

Using Trim Galore! (62), FastQ sequence files were trimmed to remove bases with quality scores less than Q30 and to discard sequences with fewer than 75 bp. To identify host-like RNA, trimmed sequences were mapped against a reference genome of *Amphiprion ocellaris* (the only publicly available clownfish genome, NCBI accession PRJNA407816) using default parameters in BBMap. The unmapped reads were retained and ribosomal reads removed using riboPicker (63). Unmapped, non-ribosomal reads were used as input for BLASTX queries against the NCBI nr database (October, 2017 release) via stand-alone BLAST version 2.6.0+. BLASTX results with bit score >50 were imported into MEGAN6 (64) with taxonomy assigned to reads using NCBI taxonomy and MEGAN6’s LCA algorithm. Functional annotation was performed in MEGAN using the SEED database. SEED is a composite, regularly curated genome annotation database (65). Gene categories that differed significantly in abundance between “fed” and “unfed” transcriptomes were identified using DESEQ2 in R, based on the raw gene count matrix exported from MEGAN6.

## DECLARATIONS

### Ethics approval and consent to participate

All live animal work was conducted in accordance with Georgia Tech IACUC protocol A15085.

### Consent for publication

Not applicable

### Availability of data and materials

The datasets generated and/or analyzed during the current study will be made publicly available in the NCBI database under BioProject ID PRJNA479844.

### Authors’ contributions

DJP and FJS conceived of the study, synthesized the results, and wrote the manuscript. DJP, MMM, FJS designed the experiments. DJP and MMM collected samples, extracted DNA and RNA, and generated sequence data. DJP analyzed the data, produced all figures, and wrote the first draft of the paper, with subsequent editing by co-authors.

## Acknowledgements

We thank John and Matthew Carberry at Sustainable Aquatics for stimulating discussion about marine aquaculture, and for their deep knowledge of clownfish husbandry in general.

## REFERENCES

1. Chung, H., Pamp, S. J., Hill, J. A., Surana, N. K., Edelman, S. M., Troy, E. B., Reading, N.C., Villablanca, E.J., Wang, S., Mora, J.R. and Umesaki, Y. Gut immune maturation depends on colonization with a host-specific microbiota. Cell, 2012;149(7):1578–1593.

2. Lee, W. J., & Hase, K.Gut microbiota–generated metabolites in animal health and disease. Nature chemical biology, 2014;10(6):416.

3. Lozupone CA, Stombaugh JI, Gordon JI, Jansson JK, Knight R. Diversity, stability and resilience of the human gut microbiota. Nature. 2012;489(7415):220–30.

4. Conlon MA, Bird AR. The impact of diet and lifestyle on gut microbiota and human health. Nutrients. 2014;7(1):17–44.

5. Galland L. The Gut Microbiome and the Brain. Journal of Medicinal Food. 2014;17(12):1261–1272.

6. Gilbert JA, Quinn RA, Debelius J, Xu ZZ, Morton J, Garg N, Jansson, J.K., Dorrestein, P.C. and Knight, R. Microbiome-wide association studies link dynamic microbial consortia to disease. Nature. 2016;535:94.

7. Thaiss, C. A., Zeevi, D., Levy, M., Zilberman-Schapira, G., Suez, J., Tengeler, A. C., Abramson L., Katz, M.N., Korem, T., Zmora, N. and Kuperman, Y. Transkingdom control of microbiota diurnal oscillations promotes metabolic homeostasis. Cell, 2014;159(3):514–529.

8. Thaiss, C. A., Levy, M., Korem, T., Dohnalová, L., Shapiro, H., Jaitin, D. A., David, E., Winter, D.R., Gury-BenAri, M., Tatirovsky, E. and Tuganbaev, T. Microbiota diurnal rhythmicity programs host transcriptome oscillations. Cell, 2016;167(6):1495–1510.

9. Voigt, R. M., Forsyth, C. B., Green, S. J., Engen, P. A., & Keshavarzian, A. Circadian rhythm and the gut microbiome. In International review of neurobiology 2016;131:193–205. Academic Press.

10. Kaczmarek, J. L., Thompson, S. V., & Holscher, H. D. Complex interactions of circadian rhythms, eating behaviors, and the gastrointestinal microbiota and their potential impact on health. Nutrition reviews, 2017;75(9):673–682.

11. Zarrinpar, A., Chaix, A., Yooseph, S., & Panda, S. Diet and feeding pattern affect the diurnal dynamics of the gut microbiome. Cell metabolism, 2014;20(6): 1006–1017.

12. Li, F., Jiang, C., Krausz, K. W., Li, Y., Albert, I., Hao, H., Fabre, K.M., Mitchell, J.B., Patterson, A.D. and Gonzalez, F.J. Microbiome remodelling leads to inhibition of intestinal farnesoid X receptor signalling and decreased obesity. Nature communications, 2013;4:2384.

13. Hehemann, J. H., Correc, G., Barbeyron, T., Helbert, W., Czjzek, M., & Michel, G. Transfer of carbohydrate-active enzymes from marine bacteria to Japanese gut microbiota. Nature, 2010;464(7290):908.

14. McNulty, N. P., Yatsunenko, T., Hsiao, A., Faith, J. J., Muegge, B. D., Goodman, A. L., Henrissat, B., Oozeer, R., Cools-Portier, S., Gobert, G. and Chervaux, C. The impact of a consortium of fermented milk strains on the gut microbiome of gnotobiotic mice and monozygotic twins. Science translational medicine, 2011;3(106):106ra106–106ra106.

15. Nayak SK. Role of gastrointestinal microbiota in fish. Aquaculture Research. 2010 Oct;41(11):1553–73.

16. Comstock, L. E., & Coyne, M. J. Bacteroides thetaiotaomicron: a dynamic, niche-adapted human symbiont. Bioessays, 2003;25(10):926–929.

17. Belenguer, A., Duncan, S. H., Calder, A. G., Holtrop, G., Louis, P., Lobley, G. E., & Flint, H. J.. 2006. Two routes of metabolic cross-feeding between Bifidobacterium adolescentis and butyrate-producing anaerobes from the human gut. Applied and environmental microbiology 72(5):3593–3599.

18. De Vuyst, L., & Leroy, F. Cross-feeding between bifidobacteria and butyrate-producing colon bacteria explains bifdobacterial competitiveness, butyrate production, and gas production. International journal of food microbiology, 2011;149(1):73–80.

19. Sung, J., Kim, S., Cabatbat, J. J. T., Jang, S., Jin, Y. S., Jung, G. Y., & Kim, P. J. Global metabolic interaction network of the human gut microbiota for context-specific community-scale analysis. Nature communications, 2017;8: 15393.

20. Van Hoek, M. J., & Merks, R. M. Emergence of microbial diversity due to cross-feeding interactions in a spatial model of gut microbial metabolism. BMC systems biology, 2017;11(1):56.

21. Espey, M. G. Role of oxygen gradients in shaping redox relationships between the human intestine and its microbiota. Free Radical Biology and Medicine, 2013; 55:130–140.

22. Albenberg, L., Esipova, T. V., Judge, C. P., Bittinger, K., Chen, J., Laughlin, A., Grunberg, S., Baldassano, R.N., Lewis, J.D., Li, H. and Thom, S.R. Correlation between intraluminal oxygen gradient and radial partitioning of intestinal microbiota. Gastroenterology 2014;147(5):1055–1063.

23. Donaldson, G. P., Lee, S. M., & Mazmanian, S. K. Gut biogeography of the bacterial microbiota. Nature Reviews Microbiology, 2016;14(1):20.

24. Gerritse, J., Schut, F., & Gottschal, J. C. Mixed chemostat cultures of obligately aerobic and fermentative or methanogenic bacteria grown under oxygen-limiting conditions. FEMS Microbiology Letters, 1990;66(1-3):87–93.

25. Johnson, K. S., & Barbehenn, R. V.Oxygen levels in the gut lumens of herbivorous insects. Journal of Insect Physiology, 2000;46(6):897–903.

26. Gross, E. M., Brune, A., & Walenciak, O. Gut pH, redox conditions and oxygen levels in an aquatic caterpillar: potential effects on the fate of ingested tannins. Journal of Insect Physiology, 2008;54(2):462–471.

27. Crawford, P. A., Crowley, J. R., Sambandam, N., Muegge, B. D., Costello, E. K., Hamady, M., Knight, R. & Gordon, J. I. Regulation of myocardial ketone body metabolism by the gut microbiota during nutrient deprivation. Proceedings of the National Academy of Sciences, 2009;106(27):11276–11281.

28. Kohl, K. D., Amaya, J., Passement, C. A., Dearing, M. D., & McCue, M. D. Unique and shared responses of the gut microbiota to prolonged fasting: a comparative study across five classes of vertebrate hosts. FEMS microbiology ecology, 2014;90(3):883–894.

29. Secor, S. M., & Carey, H. V.. Integrative physiology of fasting. Comprehensive Physiology, 2016.

30. Xing, M., Hou, Z., Yuan, J., Liu, Y., Qu, Y., & Liu, B. Taxonomic and functional metagenomic profiling of gastrointestinal tract microbiome of the farmed adult turbot (Scophthalmus maximus). FEMS microbiology ecology, 2013;86(3):432–443.

31. Givens, C. E., Ransom, B., Bano, N., & Hollibaugh, J. T. Comparison of the gut microbiomes of 12 bony fish and 3 shark species. Marine Ecology Progress Series, 2015;518:209–223. Chabrillón, M., Rico, R.M., Balebona, M.C. and Moriñigo, M.A., 2005. Adhesion to sole, Solea senegalensis Kaup, mucus of microorganisms isolated from farmed fish, and their interaction with Photobacterium damselae subsp. piscicida. Journal of Fish Diseases, 28(4), pp.229–237.

32. Chabrillón, M., Rico, R.M., Balebona, M.C. and Moriñigo, M.A., 2005. Adhesion to sole, Solea senegalensis Kaup, mucus of microorganisms isolated from farmed fish, and their interaction with Photobacterium damselae subsp. piscicida. Journal of Fish Diseases, 28(4), pp.229–237.

33. Urbanczyk, H., Ast, J. C., & Dunlap, P. V. Phylogeny, genomics, and symbiosis of Photobacterium. FEMS microbiology reviews, 2011;35(2):324–342.

34. Labella, A. M., Arahal, D. R., Castro, D., Lemos, M. L., & Borrego, J. J. Revisiting the genus Photobacterium: taxonomy, ecology and pathogenesis. Int. Microbiol, 2017;20:1–10.

35. Rastogi, R., Wu, M., DasGupta, I., & Fox, G. E. Visualization of ribosomal RNA operon copy number distribution. BMC microbiology, 2009;9(1):208.

36. Itoi, S., Okamura, T., Koyama, Y., and Sugita, H. Chitinolytic bacteria in the intestinal tract of Japanese coastal fishes. Can. J. Microbiol. 2006;52:1158–1163. doi:10.1139/w06-082

37. Ray, A. K., Ghosh, K., and Ringø, E. Enzyme-producing bacteria isolated from fish gut: a review. Aquac. Nutr. 2012;18:465–492. doi: 10.1111/j.1365-2095.2012.00943.x

38. Egerton, S., Culloty, S., Whooley, J., Stanton, C., & Ross, R. P. The gut microbiota of marine fish. Frontiers in Microbiology, 2018;9.

39. Desai, A.R., Links, M.G., Collins, S.A., Mansfield, G.S., Drew, M.D., Van Kessel, A.G. and Hill, J.E. Effects of plant-based diets on the distal gut microbiome of rainbow trout (Oncorhynchus mykiss). Aquaculture 2012;350:134–142.

40. Schmidt, V., Amaral-Zettler, L., Davidson, J., Summerfelt, S. and Good, C. Influence of fishmeal-free diets on microbial communities in Atlantic Salmon (Salmo salar) recirculation aquaculture systems. Appl Environ Microbiol 2016;82:4470–4481.

41. Turnbaugh, P. J., Ley, R. E., Mahowald, M. A., Magrini, V., Mardis, E. R., & Gordon, J. I. An obesity-associated gut microbiome with increased capacity for energy harvest. Nature, 2006;444(7122):1027.

42. Guo, X., Xia, X., Tang, R., Zhou, J., Zhao, H. and Wang, K., 2008. Development of a real-time PCR method for Firmicutes and Bacteroidetes in faeces and its application to quantify intestinal population of obese and lean pigs. Letters in applied microbiology, 47(5), pp.367–373.

43. Krajmalnik-Brown R, Ilhan ZE, Kang DW, & DiBaise JK. Effects of gut microbes on nutrient absorption and energy regulation. Nutr. Clin. Pract. 2012;27:201–14.

44. Kim, D. H., Brunt, J., & Austin, B.Microbial diversity of intestinal contents and mucus in rainbow trout (Oncorhynchus mykiss). Journal of Applied Microbiology, 2007;102(6):1654–1664.

45. Clements, K. D., Pasch, I. B., Moran, D., and Turner, S. J. Clostridia dominate 16S rRNA gene libraries prepared from the hindgut of temperate marine herbivorous fishes. Mar. Biol. 2007;150:1431–1440. doi:10.1007/s00227-006-0443-9.

46. Tailford, L. E., Owen, C. D., Walshaw, J., Crost, E. H., Hardy-Goddard, J., Le Gall, G., De Vos, W., Taylor, G., Juge, N. Discovery of intramolecular trans-sialidases in human gut microbiota suggests novel mechanisms of mucosal adaptation. Nature communications, 2015;6:7624.

47. Lopetuso, L. R., Scaldaferri, F., Petito, V., & Gasbarrini, A. Commensal Clostridia: leading players in the maintenance of gut homeostasis. Gut pathogens, 2013;5(1), 23.

48. Balcázar, J. L., De Blas, I., Ruiz-Zarzuela, I., Cunningham, D., Vendrell, D., and Muzquiz, J. L. The role of probiotics in aquaculture. Vet. Microbiol. 2006; 114:173–186. doi:10.1016/j.vetmic.2006.01.009.

49. Ramirez, R. F., and Dixon, B. A. Enzyme production by obligate intestinal anaerobic bacteria isolated from oscars (Astronotus ocellatus), angelfish (Pterophyllum scalare) and southern flounder (Paralichthys lethostigma). Aquaculture, 2003;227:417–426. doi:10.1016/S0044-8486(03)00520-9.

50. Matches J R, Liston J, Curran D. Clostridium perfringens in the environment. Appl Microbiol. 1974;28:655–660.

51. Sabry M, Abd El-Moein K, Hamza E, Abdel Kader F. Occurrence of Clostridium perfringens Types A, E, and C in Fresh Fish and Its Public Health Significance. J Food Prot. 2016;79(6):994–1000. doi: 10.4315/0362-028X.JFP-15-569. PubMed PMID: 27296604.

52. Deplancke B, Vidal O, Ganessunker D, Donovan SM, Mackie RI, Gaskins HR. Selective growth of mucolytic bacteria including Clostridium perfringens in a neonatal piglet model of total parenteral nutrition. Am J Clin Nutr. 2002 Nov;76(5):1117–25.

53. Machado H, & Gram L. Comparative Genomics Reveals High Genomic Diversity in the Genus Photobacterium. Frontiers in Microbiology. 2017;8:1204. doi:10.3389/fmicb.2017.01204.

54. Gu J, Neary J, Cai, H., Moshfeghian, A., Rodriguez, S.A., Lilburn, T.G. and Wang, Y. Genomic and systems evolution in Vibrionaceae species. BMC Genomics. 2009;10(Suppl 1):S11. doi:10.1186/1471-2164-10-S1-S11.

55. Lin H, Yu M, Wang X, Zhang X-H. Comparative genomic analysis reveals the evolution and environmental adaptation strategies of vibrios. BMC Genomics. 2018;19:135. doi:10.1186/s12864-018-4531-2.

56. Caporaso, J. G., Lauber, C. L., Walters, W. A., Berg-Lyons, D., Lozupone, C. A., Turnbaugh, P. J., & Knight, R. Global patterns of 16S rRNA diversity at a depth of millions of sequences per sample. Proceedings of the National Academy of Sciences, 2011;108(Supplement 1):4516–4522.

57. Kozich, J. J., Westcott, S. L., Baxter, N. T., Highlander, S. K., & Schloss, P. D. Development of a dual-index sequencing strategy and curation pipeline for analyzing amplicon sequence data on the MiSeq Illumina sequencing platform. Applied and environmental microbiology, 2013;79(17):5112–5120.

58. Amir, A., McDonald, D., Navas-Molina, J. A., Kopylova, E., Morton, J. T., Xu, Z. Z.,& Knight, R. 2017. Deblur rapidly resolves single-nucleotide community sequence patterns. MSystems, 2(2):e00191–16.

59. Mandal, S., Van Treuren, W., White, R. A., Eggesbø, M., Knight, R., & Peddada, S. D.. Analysis of composition of microbiomes: a novel method for studying microbial composition. Microbial ecology in health and disease, 2015;26(1): 27663.

60. Langille, M. G., Zaneveld, J., Caporaso, J. G., McDonald, D., Knights, D., Reyes, J. A., Clemente, J.C., Burkepile, D.E., Thurber, R.L.V., Knight, R. and Beiko, R.G. Predictive functional profiling of microbial communities using 16S rRNA marker gene sequences. Nature biotechnology, 2013;31(9): 814.

61. Ling, K. M., & Ghaffar, M. A. Estimation of gastric emptying time (GET) in clownfish (Amphiprion ocellaris) using X-radiography technique. In AIP Conference Proceedings 2014;1614(1):624–628.

62. Krueger, F.,. Trim galore. A wrapper tool around Cutadapt and FastQC to consistently apply quality and adapter trimming to FastQ files. 2015.

63. Schmieder, R., Lim, Y. W., & Edwards, R.. Identification and removal of ribosomal RNA sequences from metatranscriptomes. Bioinformatics, 2011; 28(3):433–435.

64. Huson, D. H., Beier, S., Flade, I., Gorska, A., El-Hadidi, M., Mitra,S., & Tappu, R. MEGAN community edition-interactive exploration and analysis of large-scale microbiome sequencing data. PLoS computational biology, 2016;12(6):e1004957.

65. Overbeek, R.F., Begley, T., Butler, R.M., Choudhuri, J.V., Chuang, H.Y., Cohoon, M., de Crecy-Lagard, V., Diaz, N., Disz, T., Edwards, R., and Fonstein, M. The subsystems approach to genome annotation and its use in the project to annotate 1000 genomes. Nucleic Acids Res,. 2005;33:5691–5702.

